# Independent Theta Phase Coding Accounts for CA1 Population Sequences and Enables Flexible Remapping

**DOI:** 10.1101/005066

**Authors:** Angus Chadwick, Mark C. W. van Rossum, Matthew F. Nolan

## Abstract

Populations of hippocampal place cells encode an animal’s past, current and future location through sequences of action potentials generated within each cycle of the network theta rhythm. These sequential representations have been suggested to result from temporally coordinated synaptic interactions within and between cell assemblies. In contrast, we show that a model based on rate and phase coding in independent neurons is sufficient to explain the organization of CA1 population activity during theta states. We show that CA1 population activity can be described as an evolving traveling wave that exhibits phase coding, rate coding, spike sequences and that generates an emergent population theta rhythm. We identify measures of global remapping and intracellular theta dynamics as critical for distinguishing mechanisms for pacemaking and coordination of sequential population activity. Our analysis suggests that independent coding enables flexible generation of sequential population activity within the duration of a single theta cycle.

## INTRODUCTION

Cognitive processes are thought to involve the organization of neuronal activity into phase sequences, reflecting sequential activation of different cell assemblies (Hebb, 1949; Harris, 2005). During navigation, populations of place cells in the CA1 region of the hippocampus generate phase sequences structured relative to the theta rhythm (e.g., Skaggs et al., 1996; Foster and Wilson, 2007). As an animal moves through the firing field of a single CA1 neuron, there is an advance in the phase of its action potentials relative to the extracellular theta cycle (O’Keefe and Recce, 1993). Thus, populations of CA1 neurons active at a particular phase of theta encode the animal’s recent, current, or future positions (Figure 1A, B). One explanation for these observations is that synaptic output from an active cell assembly ensures its other members are synchronously activated and in addition drives subsequent activation of different assemblies to generate a phase sequence (Figure 1C) (Harris, 2005). We refer to this as the *coordinated assembly hypothesis*. An alternative possibility is that single cell coding is sufficient to account for population activity. According to this hypothesis, currently active assemblies do not determine the identity of future assemblies (Figure 1D). We refer to this as the *independent coding hypothesis*.

**Figure 1:**
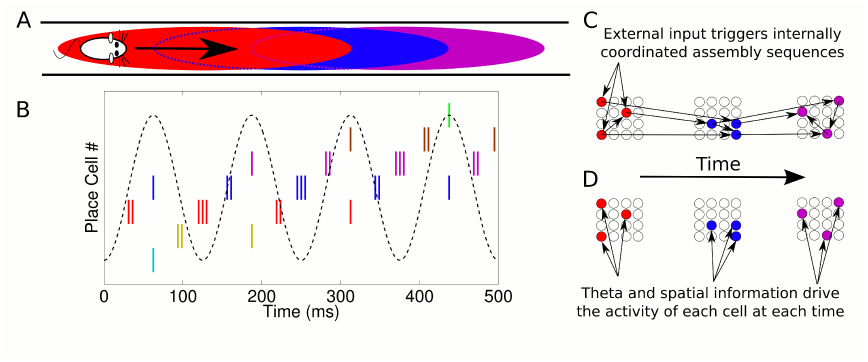
Phase sequences in a place cell population. (A) During navigation, place cells are sequentially activated along a route. (B) Within each theta cycle, this slow behavioral sequence of place cell activations is played out on a compressed timescale as a theta sequence. Theta sequences involve both rate and phase modulation of individual cells, but it remains unclear whether additional coordination between cells is present. (C) Internal coordination may bind CA1 cells into assemblies, and sequential assemblies may be chained together synaptically. This would require specific inter- and intra-assembly patterns of synaptic connectivity within the network. (D) Alternatively, according to the independent coding hypothesis, each cell could be governed by theta phase precession without additional coordination.

Since these coding schemes lead to different views on the nature of the information transferred from hippocampus to neocortex and on the role of CA1 during theta states, it is important to distinguish between them. While considerable experimental evidence has been suggested to support the coordinated coding hypothesis (e.g., Harris et al., 2003; Foster and Wilson, 2007; Maurer et al., 2011; Gupta et al., 2012), the extent to which complex sequences of activity in large neuronal populations can be accounted for by independent coding is not clear. To address this we developed a phenomenological model of place cell activity during navigation. This model is based on rate coding across a place field and phase precession against a fixed theta rhythm. The independent coding hypothesis predicts that this model, when generalized to a population of independent cells, will be sufficient to explain the spatiotemporal dynamics of cell assemblies in CA1. In contrast, the coordinated assembly hypothesis predicts that groups of cells show additional coordination beyond that imposed by a fixed firing rate and phase code (Harris et al., 2003; Harris, 2005). In this case, independent coding would not be sufficient to explain the detailed dynamics of CA1 cell assemblies. In the independent coding model that we develop, phase coding generates a traveling wave which propagates through the population to form spike sequences. This wave is constrained by a slower moving modulatory envelope which generates spatially localized place fields. Importantly, this model replicates experimental data previously interpreted as evidence for the coordinated assembly hypothesis (Harris et al., 2003; Foster and Wilson, 2007; Maurer et al., 2011; Gupta et al., 2012), despite the absence of coordination within or between assemblies. We show how the independent coding model leads to new and experimentally testable predictions for membrane potential oscillations and place field remapping that distinguish circuit mechanisms underlying theta sequences. We demonstrate that a key advantage of independent coding by CA1 neurons is to allow flexible global remapping of population activity while maintaining the ability to generate coherent theta sequences in multiple environments.

## RESULTS

### Single Cell Coding Model

To test the independent coding hypothesis, we developed a phenomenological model which generates spiking activity for a single place cell during navigation. We modeled the firing rate field using a Gaussian tuning curve:

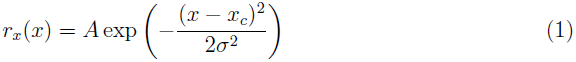

where *r_x_* describes firing rate when the animal is at location *x* within a place field with center *x_c_*, width *σ* and maximum rate *A* (Figure 2A, top panel). Simultaneously, we modeled the firing phase using a circular Gaussian:

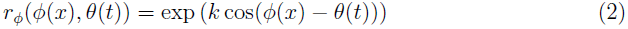

where *r_ϕ_* describes the firing probability of the neuron at each theta phase at a given location (Figure 2B). Here, *θ*(*t*) = 2*πf*_*θ*_*t* is the LFP theta phase at time *t* and *ϕ*(*x*) is the preferred firing phase associated with the animal’s location *x*, denoted the *encoded phase*. The encoded phase *ϕ*(*x*) is defined to precess linearly across the place field (Figure 2A, bottom panel; Appendix: A1). The *phase locking* parameter *k* determines the precision at which the encoded phase is represented in the spike output (Figure 2B). The total activity of the cell is given by the product of these two components *r* = *r_x_r_ϕ_*. The phase locking can be set so that the cell exhibits only rate coding (at *k* = 0, where *r* = *r_x_*), only phase coding (as *k* → ∞, where all spikes occur at exactly the encoded phase *ϕ*(*x*)) or anywhere in between (Figure 2C).

**Figure 2:**
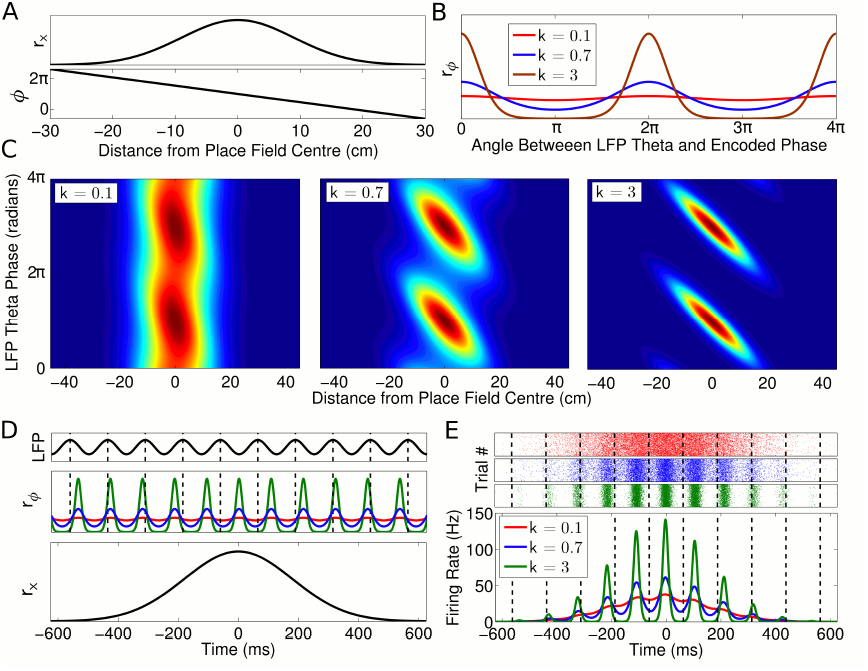
Single cell coding model. (A) Firing rate and phase at different locations within a cell’s place field are determined by a Gaussian tuning curve *r_x_* and linearly precessing encoded phase *ϕ* respectively. (B) The dependence of single cell activity on the LFP theta phase *θ* is modeled by a second tuning curve *r_ϕ_* which depends on the angle between the LFP theta phase *θ* and encoded phase *ϕ* at the animal’s location. The phase locking parameter *k* controls the precision of the phase code. (C) The combined dependence of single cell activity on location and LFP theta phase. (D) Temporal evolution of the rate and phase tuning curves for a single cell as a rat passes through the place field at constant speed. (E) The total firing rate corresponding to (D), and spiking activity on 1000 identical runs.

To model place cell activity during navigation on a linear track, we set *x*(*t*) = *vt*, where *v* is the running speed (Figure 2D, E). This causes the encoded phase *ϕ*(*t*) to precess linearly in time at a rate *f_ϕ_* which is directly proportional to running speed and inversely proportional to place field size, as in experimental data (Huxter et al., 2003; Geisler et al., 2007). To generate spikes we used an inhomogeneous Poisson process with an instantaneous rate *r* = *r_x_r_ϕ_*. We normalized the firing rate such that the average number of spikes fired on a pass through a place field is independent of running speed (see Appendix: A2) (Huxter et al., 2003). If the phase *ϕ*(*x*) at each location in the place field is fixed, the full rate and phase coding properties of a cell are encompassed by three independent parameters - the width of the spatial tuning curve *σ*, the degree of phase locking *k* and the average number of spikes per pass *N*_spikes_. We find that in this model phase precession (Figure 2C) and firing rate modulation as a function of time (Figure 2E) closely resemble experimental observations (e.g., Skaggs et al., 1996; Mizuseki and Buzsaki, 2014).

Place cells often show variations in firing rate in response to nonspatial factors relevant to a particular task (e.g., Wood et al., 2000; Griffin et al., 2007; Fyhn et al., 2007; Allen et al., 2012). In our model, such multiplexing of additional rate coded information can be achieved by varying the number of spikes per pass *N*_spikes_ without interfering with the other parameters *ϕ*(*x*), *σ* and *k* (Figure S1).

### Independent phase coding generates traveling waves

Given this single cell model and assuming an independent population code, we asked how CA1 population activity evolves during navigation. To map the spatiotemporal dynamics of the population activity onto the physical space navigated by the animal, we analyzed the distributions of the rate components *r_x_* and phase components *r_ϕ_* of activity in cell populations organized according to the location *x_c_* of each place field (Appendix: A3).

Our model naturally separates population activity into two components at different timescales: the slow behavioral timescale at which the rat navigates through space and a fast theta timescale at which trajectories are compressed into theta sequences. While the rat moves through the environment, the spatial tuning curves *r_x_*(*x*) generate a slow moving ‘bump’ of activity which, by definition, is comoving with the rat (Figure 3A top, black). Simultaneously, the phasic component *r_ϕ_*(*ϕ*(*x*)*, θ*(*t*)) instantiates a traveling wave (Figure 3A top, red). Due to the precession of *ϕ*(*t*), the wave propagates forward through the network at a speed faster than the bump, resulting in sequential activation of cells along a trajectory on a compressed timescale. The slower bump of activity acts as an envelope for the traveling wave, limiting its spatial extent to one place field (Figure 3A bottom). The continuous forward movement of the traveling wave is translated into discrete, repeating theta sequences via a shifting phase relationship to the slow moving component (Figure 3B-D, Supplemental Video 1). Moreover, this shifting phase relationship generates global theta oscillations at exactly the LFP frequency that cells were defined to precess against (Figure 3B top panel). Thus, our model can be recast in terms of the dynamics of a propagating wavepacket comprising two components, with network theta resulting from their interaction. While we originally defined single cells to precess against a predetermined “pacemaker” theta rhythm, when the model is recast in this form the same theta oscillation instead emerges independently from the population. A parsimonious interpretation is that the traveling wave dynamics outlined here could generate both the LFP theta rhythm and the precession of cells against it, without an external pacemaker.

**Figure 3:**
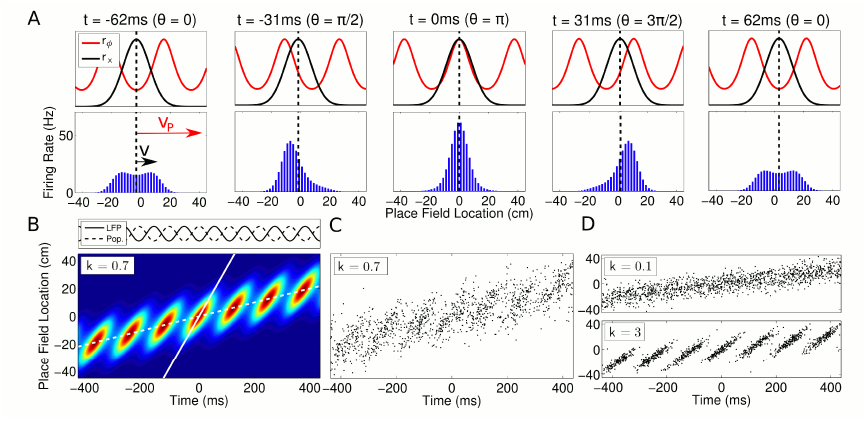
Spatiotemporal dynamics of CA1 populations. (A) *Top:* Population dynamics during a single theta cycle on a linear track after ordering cells according to their place field center *x_c_* in physical space. The two components of the population activity are shown - the slow moving envelope (black) and the fast moving traveling wave (red), which give rise to rate coding and phase coding respectively (cf. Figure 2). *Bottom:* Resulting firing rates across the population. When the traveling wave and envelope are aligned, the population activity is highest (middle panel). The dashed line shows the location of the rat at each instant. **B:** Firing rate in the population over seven consecutive theta cycles. The fast and slow slopes are shown (solid and dashed lines respectively), corresponding to the speeds of the traveling wave and envelope as shown in part **A**. The top panel shows the LFP theta oscillations and emergent population theta oscillations, which are generated by the changing population activity as the traveling wave shifts in phase relative to the slower envelope (see Supplemental Video 1). (C, D) The spiking activity for a population of 180 cells. All panels used *v* = 50 cm/s, so that *v_p_* = 350 cm/s and *c* = 7.

While the emergence of global theta oscillations in networks of faster oscillating place cells has been identified previously (Geisler et al., 2010), that work assumed a fixed running speed and fixed, experimentally determined, temporal delays between cells. In contrast, our model based on single cell coding principles allows an analysis in which only place field configurations and navigational trajectories are required to fully predict the population dynamics at any running speed. The spiking delays between cells in our model are determined by speed of the fast moving traveling wave *v_p_*, which is related to the rat’s running speed *v* by:

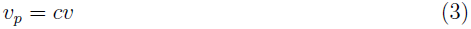

where *c* is called the *compression factor*. This factor is equivalent to the ratio of the rat’s actual velocity and the velocity of the representation within a theta cycle and has been quantified in previous experimental work (Skaggs et al., 1996; Dragoi and Buzsáki, 2006; Geisler et al., 2007; Maurer et al., 2011), although the relationship to the traveling wave model developed here was not previously identified (see Appendix: A2 for derivation).

A novel finding of our model is that for an independent population code the compression factor naturally depends on running speed. This change in compression factor with running speed allows the network to maintain a fixed population frequency while running speed and single unit frequency vary:

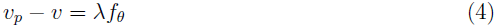

where *λ* is the wavelength of the traveling wave (equal to the size of a place field, measured as the distance over which a full cycle of phase is precessed (Maurer et al., 2006)) and *v_p_* − *v* is held constant across running speeds by the changing compression factor.

### Independent coding accounts for apparent peer-dependence of CA1 activity

Having derived a model for population activity based on independent coding, we next asked if it accounts for observations previously interpreted to imply coordination within and between assemblies (Harris et al., 2003; Foster and Wilson, 2007; Maurer et al., 2011; Gupta et al., 2012). We show below that these observations can be accounted for by independent neurons using the traveling wave model.

We first assessed whether independent coding accounts for membership of cell assemblies. The predictability of a single place cell’s activity appears to improve when, in addition to information about LFP theta phase and spatial location, the activity of its peer cells is also taken into account, with coordination between cells at the gamma timescale being implicated (Harris et al., 2003). Because this improved predictability directly implies interactions between CA1 neurons, it constitutes strong evidence against the independent coding hypothesis. However, the prediction analysis of Harris et al. (2003) assumes that firing phase is independent of movement direction in an open environment. In contrast, more recent experimental data show that in open environments firing phase always precesses from late to early phases of theta, so that firing phase at a specific location depends on the direction of travel (Huxter et al., 2008; Climer et al., 2013; Jeewajee et al., 2014). Therefore, to test if the apparent peer dependence of place cell activity is in fact consistent with independent coding, we extended the traveling wave model to account for phase precession in open environments (Appendix: A5). We then constructed *phase fields* following the approach of (Harris et al., 2003), in which firing phase is averaged over all running directions and separately constructed directional phase fields consistent with recent experimental observations (Huxter et al., 2008; Climer et al., 2013; Jeewajee et al., 2014). We then calculated the predictability of neuronal firing patterns generated by the independent coding model. For simplicity, we considered the problem in one dimension, treating separately passes from right to left, left to right and the combined data in order to generate the directional and nondirectional phase fields (Figure 4A&B respectively).

**Figure 4:**
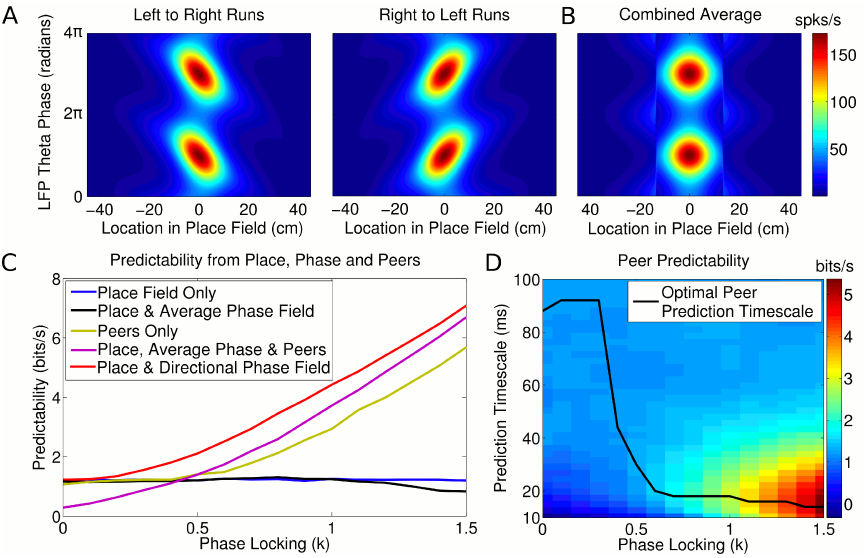
Peer prediction analysis for an independent population code. (A) Combined place and phase fields constructed from simulated data using only runs with a single direction. (B) Place/phase field constructed from a combination of both running directions, as used by Harris et al. (2003). (C) Predictability analysis, using various combinations of place, phase and peer activity. When using the nondirectional phase field of Harris et al. (2003), an additional peer predictability emerges (black vs green and purple). However, this additional predictability is seen to be illusory if the directional phase field is used to predict activity (red). (D) Dependence of peer predictability on the peer prediction timescale and phase locking of individual cells. The heat map shows the predictability of a cell’s activity from peer activity (cf. part C, green line). The optimal peer prediction timescale depends on the amount of phase locking, showing that the characteristic timescale of peer correlations reflects independent phase precession of single cells rather than transient gamma synchronization of cell assemblies.

We find that nondirectional phase fields (Figure 4B), as assumed by (Harris et al., 2003), yield little improvement in predictability of a neuron’s firing compared with predictions based on the place field alone, and for high phase locking are detrimental (Figure 4C, blue vs black). Consistent with the findings of Harris et al. (2003), peer prediction provides a higher level of information about a neuron’s firing than predictions based on place and nondirectional phase fields, despite the absence of intra-assembly coordination (Figure 4C, green and purple). However, peer prediction is unable to improve upon predictability based on place fields and directional phase fields (Figure 4C, red). Therefore, previous evidence for intra-assembly coordination can be explained by a failure to account for the phase dependence of CA1 firing. Instead, our analysis indicates that independent phase precession of CA1 neurons is sufficient to account for membership of CA1 assemblies.

Because peers share a relationship to a common theta activity and implement similar rules for generation of firing, a cell’s activity in the independent coding model can nevertheless be predicted from that of its peers in the absence of information about location or theta phase (Figure 4C, green). The quality of this prediction is dependent on the timescale at which peer activity is included in the analysis, so that the optimal timescale for peer prediction provides a measure of the temporal resolution of assembly formation. In experimental data the optimal timescale for peer prediction is approximately 20 ms, which corresponds to that of the gamma rhythm and the membrane time constant of CA1 neurons (Harris et al., 2003). We find that in the independent coding model the optimal peer prediction timescale depends strongly on phase locking (Figure 4D). Even though the model does not incorporate gamma oscillations or neuronal membrane properties, high values of phase locking also show a striking peak in peer predictability around the 20 ms range (Figure 4D). We show below that for running speeds in the range 35 *-* 75 cm/s phase locking is likely to lie within the range at which the observed 20 ms prediction timescale dominates. Thus, the 20 ms timescales found both here and experimentally are explainable as a signature of the common, independent phase locking of place cells to the theta rhythm, rather than transient gamma coordination or intrinsic properties of CA1 neurons.

### Independent coding accounts for phase sequences

We next asked if independent coding can account for the sequences of spiking activity observed in recordings of CA1 place cell populations (Foster and Wilson, 2007; Maurer et al., 2011; Gupta et al., 2012). We focus initially on the path length encoded by spike sequences, which we define as the length of trajectory represented by the sequence of spikes within a single theta cycle. Experimental data show that this path length varies with running speed (Maurer et al., 2011; Gupta et al., 2012), but it is not clear whether this phenomenon is a feature of independent coding or instead results from coordination between assemblies. We derived analytical approximations to the sequence path length for strong phase coding, where *k → ∞* (Appendix: A2) which predict a linear increase in sequence path length with running speed, but with a lower gradient than that found experimentally (Maurer et al., 2011). Hence, strong phase coding does not quantitatively explain the changes in sequence properties with running speed.

We reasoned that independent coding might still explain observed sequence path lengths if a more realistic tradeoff between rate and phase coding is taken into account. To test this, we varied phase locking *k* and decoded the path length following the method of Maurer et al. (2011), which decodes the location represented by the population at each time bin in a theta cycle to estimate the encoded trajectory. We found that a good match to the data of Maurer et al. (2011) requires that the degree of phase locking increases with running speed (Figure 5A). This is due to the dependence of the decoded path length on the amount of phase locking (Figure S2A).

**Figure 5:**
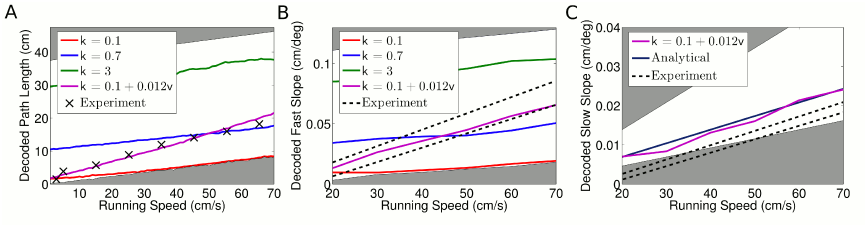
Decoded sequence path lengths and population activity propagation speeds. (A) The decoded path length increases linearly with running speed, but a dependence of phase locking on running speed is required to account for the experimental data. The shaded regions show upper and lower bounds (*k* = 0 and *k* = *∞*). (B) Dependence of decoded fast slope on running speed (cf. our Figure 3B; Maurer et al. (2011) Figure 3). Again, a match to the data requires a velocity dependent phase locking. (C) The slow slope was accurately decoded to the analytical value, where the population travels at the running speed *v*. Bounds show LFP theta frequencies below 4 Hz (upper bound) and above 12 Hz (lower bound).

To further test the traveling wave model, we analyzed the fast and slow components of the movement of the activity bump (*v* and *v_p_*). Following again the methods used by Maurer et al. (2011) to decode the fast and slow slopes shown in Figure 5B from spiking data, we found that the dependence of the decoded fast slope on running speed in our simulated data matches experimental data when phase locking is made dependent on running speed (Figure 5B, S2B). Hence, both the decoded sequence path length and theta-compressed propagation speed in the traveling wave model match experimental data provided the temporal resolution of spike output increases linearly with running speed. This dependence of *k* on running speed is consistent with the observed increase in LFP theta amplitude (McFarland et al., 1975; Maurer et al., 2005; Patel et al., 2012), and is a novel prediction made by our model.

We also asked whether the independent coding model can account for experimental measurements indicating that the compression factor scales inversely with running speed (Maurer et al., 2011). Having accurately reproduced the fast slope only the slow slope (which represents the overall movement of the population activity on a behavioral timescale) is required to reproduce the reported compression factor. However, while our decoded values for the slow slope closely matched the true value based on the rat’s running speed, the values reported by Maurer et al. (2011) are considerably lower (Figure 5C) and if correct would suggest that the population consistently moved more slowly than the rat, even moving backwards while the animal remained still. This inconsistency precluded a comparison to the compression factor in our model.

Independent experimental support for the notion of inter-assembly coordination comes from an analysis suggesting that single cell phase precession is less precise than observed theta sequences (Foster and Wilson, 2007). This conclusion relies on a shuffling analysis which preserves the statistics of single cell phase precession yet reduces intra-sequence correlations. However, performing the same shuffling analysis on data generated by our independent coding model reproduced this result (t-test, *p* < 10^−46^) (Figure S3). The effect arises because the shuffling procedure does not preserve the temporal structure of single cell phase precession, despite preserving the phasic structure (Figure S3A, B). Hence, the phase-position correlations are unaffected, while the time-position correlations and hence sequence correlations are disrupted (Figure S3C, D). Thus, these experimental observations can be accounted for by independent coding.

Finally, precise coordination of theta sequences has been suggested on the basis that theta sequence properties vary according to environmental features such as landmarks and behavioral factors such as acceleration (Gupta et al., 2012). To establish whether independent coding could also account for these results, we generated data from our traveling wave model and applied the sequence identification and decoding analyses reported by Gupta et al. (2012). We found that, even for simulated data based on pure rate coding with no theta modulation (*k* = 0), this analysis detected a fraction of “significant theta sequences” similar to that reported for experimental data (Figure 6A, B). Moreover, applying the reported Bayesian decoding analysis yielded similar decoded path lengths to those found experimentally, despite the absence of theta sequences in the simulated data (Figure 6C). Since these results can be reproduced with data containing no sequences at all, they lack the specificity required to analyze the trajectories represented by theta sequences. Instead, as the results are reproducible with purely rate coded activity, they are consistent with variations in rate coding across behavioral settings (e.g., Wood et al., 2000; Fyhn et al., 2007; Allen et al., 2012).

**Figure 6:**
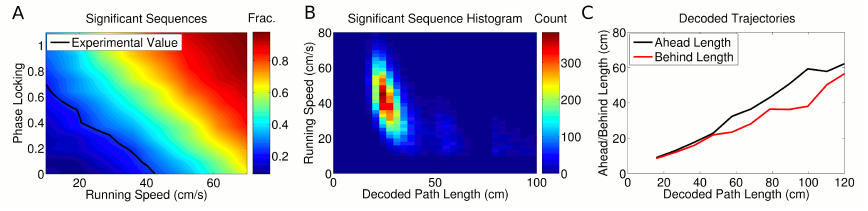
Analysis of individual sequence statistics. (A) The fraction of “significant sequences” generated under independent coding according to the Gupta et al. (2012) analysis, as a function of running speed and phase locking (for simulated data). Large fractions of significant sequences are generated even without phase coding or theta sequences within the assembly (i.e. at *k* = 0). The black line shows the fraction reported experimentally. (B) and (C) used only rate coding (*k* = 0), and therefore do not contain any theta sequences above chance. (B) The distribution of significant sequences over running speed and decoded path length, as calculated by Gupta et al. (2012) (cf. their Fig 1c). (C) The relationship between decoded path length and decoded ahead and behind lengths for significant sequences, using the same simulated data as the previous panel. This replicates the experimental data (cf. Fig 4a, b in Gupta et al. (2012)), showing that similar path lengths are decoded by this algorithm even when theta sequences are not present.

In total, our analysis demonstrates that a traveling wave model based on independent phase coding for CA1 theta states is consistent with existing experimental data. Thus, neither intra- nor inter-assembly interactions are required to explain CA1 activity during theta states. We show that the spatiotemporal organization of CA1 population activity within theta oscillations can be explained by independent phase precession against an externally fixed pacemaker rhythm. Alternatively, such populations could instead generate their own theta frequency network oscillations, so that CA1 theta activity can also be explained in terms of population coding with emergent theta frequency dynamics.

### Linear phase coding constrains global remapping

When an animal is moved between environments, the relative locations at which place cells in CA1 fire remap independently of one another (e.g., O’Keefe and Conway, 1978; Wilson and McNaughton, 1993). Due to the limited storage capacity for temporal sequences in neural networks (Sompolinsky and Kanter, 1986; Kleinfeld, 1986; Reiss and Taylor, 1991; Bressloff and Taylor, 1999; Leibold and Kempter, 2006), this global remapping of spatial representations poses a challenge for generation of theta sequences through coordinated assemblies as synaptic interactions that promote formation of sequences in one environment would be expected to interfere with sequences in a second environment. In comparison, it is not yet clear whether the independent coding model faces similar constraints on sequence generation across different spatial representations. We therefore addressed the feasibility of maintaining theta sequences following remapping given the assumptions that underpin our independent coding model.

We first consider the possibility that following remapping the phase lags between cell pairs remain fixed - that is, while two cells may be assigned new firing rate fields, their relative spike timing within a theta cycle does not change. This scenario would occur if the phase lags associated with linear phase precession were generated by intrinsic network architectures (e.g., Diba and Buzsáki, 2008; Dragoi and Tonegawa, 2011, 2013) or upstream pacemaker inputs. To illustrate the consequences of such mechanisms for global remapping, we first consider representation of a linear track by a population of place cells with distinct rate and phase fields (Figure 7A). Our description of phase precession implies a mechanism to establish a cell’s phase field. With linear phase coding a cell’s phase is maintained outside of its firing field such that the phase lag between two neurons depends linearly on the distance between their place field centers, while cells separated by multiples of a place field width share the same phase (Figure 7A). Each cell pair therefore has a fixed phase lag and all cells can in principle be mapped onto a single chart describing their phase ordering. If this mechanism for determining phase ordering is hardwired, then following arbitrary global remapping, cells with nearby place field locations will in most cases no longer share similar phases (Figure 7B). As a result, theta sequences and the global population theta will in general be abolished (Figure 7B). However, there exist a limited set of remappings which in this scenario do not disrupt the sequential structure of the population (e.g., Figure 7C). On a linear track, these remappings are: translation of all place fields by a fixed amount, scaling of all place fields by a fixed amount and permuting the place field locations of any cell pair with zero phase lag.

**Figure 7:**
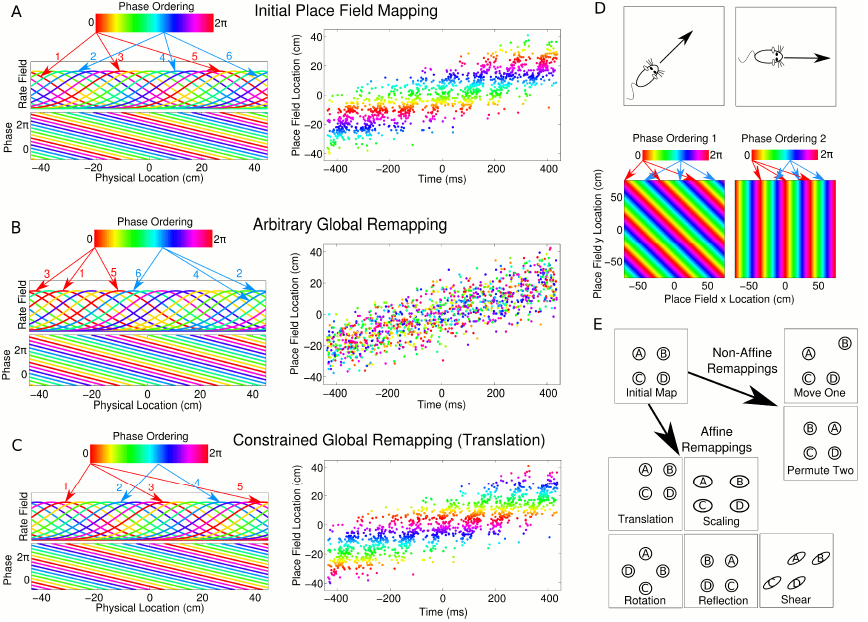
Properties of CA1 populations governed by linear phase coding. (A) On a linear track, cells which precess linearly in phase maintain fixed delays. This is illustrated as a phase ordering (colored bar), which describes the relative phase of each cell (arrows show locations of cells at each phase). Each cell has a constant, running speed dependent frequency and a fixed phase offset to each other cell. (B) A complete global remapping with phase lags between cells held fixed. Theta sequences and population oscillations are abolished. (C) In a constrained place field remapping, theta sequences are preserved. (D) In open environments, phase lags depend on running direction. The set of population phase lag configurations needed to generate sequences in each direction is called a phase chart. (E) If a population has a fixed phase chart, the possible remappings are restricted to affine transformations.

When considering global remapping in an open environment similar constraints apply. Because the phase lag between any two cells depends on running direction (e.g., Huxter et al., 2008), the population phase ordering must always be aligned with the direction of movement (Figure 7D). Hence, in open environments, the notion of a phase chart must be extended to include a fixed phase ordering for each running direction. Given such a fixed phase chart, a set of remappings known as *affine transformations* preserve the correct theta dynamics (see Appendix: A6). Such remappings consist of combinations of linear transformations (scaling, shear, rotation and reflection) and translations (Figure 7E). Remappings based on permutation of place field locations of synchronous cells, which are permissible in one dimensional environments, are no longer tenable in the two dimensional case due to constraints over each running direction.

Since place cell ensembles support statistically complete (i.e. non-affine) remappings (e.g., O’Keefe and Conway, 1978), CA1 network dynamics are not consistent with the model outlined above. Nevertheless, it remains possible that CA1 theta dynamics are based on fixed phase charts, provided that multiple such phase charts are available to the network. In this case, each complete remapping recruits a different phase chart, fixing a new set of phase lags. The number of possible global remappings that maintain theta sequences is then determined by the number of available phase charts. Such a possibility is consistent with recent suggestions of fixed sequential architectures (Dragoi and Tonegawa, 2011, 2013) and has not been ruled out in CA1. It is also of interest that affine transformations are consistent with the observed remapping properties in grid modules (Fyhn et al., 2007), suggesting that a single phase chart might be associated with a grid module.

### Sigmoidal phase coding enables theta sequence generation and flexible global remapping

Is it possible to overcome the constraints imposed on remapping by fixed phase charts in independent coding models? We reasoned that experimental data on phase precession only imply that phase precesses within a cell’s firing field and need not constrain a cell’s phase outside of its firing field. We therefore implemented a version of the independent coding model in which firing phase has a sigmoidal relationship with location (Figure 8A-B, solid line), so that phase precesses within the firing field but not outside of the field. In this case, each cell’s intrinsic frequency is increased within the spatial firing field, whereas outside the firing field it has the same frequency as the population theta rhythm (Figure 8C, solid line). This is in contrast to the linear phase model in which each cell’s intrinsic frequency is always faster than the population oscillation (Figure 8C, dashed line). When spiking activity in a population of cells with sigmoidal phase coding is considered (Figure 8D-F), phase precession and sequence generation are similar to models in which cells have linear phase coding. However, in addition, sigmoidal phase coding enables theta sequences to be generated after any arbitrary global remapping (Figure 8G). This flexible global remapping is in contrast to the scrambling of theta sequences that typically occurs with global remapping when cells have linear phase coding (Figure 8G).

**Figure 8:**
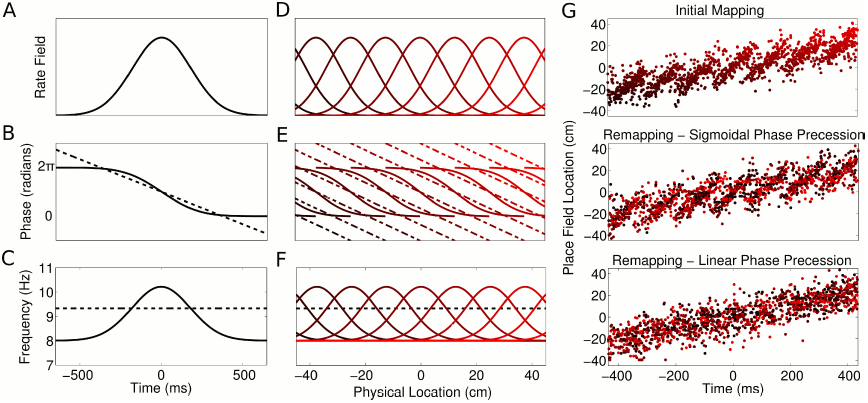
Properties of CA1 populations governed by sigmoidal phase coding. (A-C) Firing rate and intracellular phase and frequency in the linear (dashed lines) and sigmoidal models (solid lines) during the crossing of a place field. In the sigmoidal model, phase precession is initiated inside the place field by an elevation of intracellular frequency from baseline. (D-F) Firing rate and intracellular phase and frequency for a place cell population on a linear track. In the sigmoidal model, an intracellular theta phase between cell pairs develops as the animal moves through their place fields. Outside their place fields, cell pairs are synchronized. (G) Global remapping in the linear and sigmoidal models. In the sigmoidal model, the intracellular dynamics allow arbitrary remapping without disrupting population sequences.

Linear and sigmoidal models of phase coding lead to distinct experimentally testable predictions. Recordings of membrane potentials of CA1 neurons in behaving animals show that spikes precess against the LFP but always occur around the peak of a cells intrinsic membrane potential oscillation (MPO) (Harvey et al., 2009). Therefore the intrinsic phase of each cell in our model (Figure 2D, E) can be considered equivalent to MPO phase. While data concerning the MPO phase outside of the firing field are limited, this will likely distinguish generation of theta sequences based on a linear and sigmoidal phase coding. If CA1 implements linear phase coding, then the MPO of each cell should precess linearly in time against LFP theta at a fixed (velocity dependent) frequency, both when the animal is inside the place field and when the animal is at locations where the cell is silent (Figure 8A-C, dashed line). Alternatively, sigmoidal phase coding predicts that precession of the MPO against the LFP occurs only inside the firing rate field (Figure 8A, B solid line) and that the MPO drops back to the LFP frequency outside of the place field (Figure 8C solid line). A further prediction of sigmoidal coding is that the phase lag between any two cells changes when the animal moves through their place fields, whereas outside their place fields the cells are synchronized with each other and with the LFP (Supplementary Video 2). Finally, phase precession under the sigmoidal model behaves differently to the linear model in open environments. In the linear model, the phase chart fixes a different population phase ordering for each running direction, so that spike phase depends on the location of the animal and the instantaneous direction of movement. In the sigmoidal model, however, each cell has a location dependent frequency, so that the spike phase depends on the complete trajectory through the place field and no explicit directional information is required (see Appendix: A5). In summary, evaluation of theta sequences following global remapping and of theta phase within and outside of a cells firing field will be critical for distinguishing models of theta generation.

## DISCUSSION

Our model of the spiking activity of CA1 populations governed by independent phase coding demonstrates how complex and highly structured population sequences can be generated with minimal coordination between neurons. In contrast to previous suggestions (Harris et al., 2003; Foster and Wilson, 2007; Maurer et al., 2011; Gupta et al., 2012), we found that the population activity observed in CA1 can be accounted for by phase precession in independent cells, without interactions within or between cell assemblies. The independent coding hypothesis leads to a novel view of the CA1 population as a fast moving traveling wave with a slower modulatory envelope. This model exhibits rate coding, global theta oscillations and phase precession against this self generated rhythm. Amplitude modulation of the envelope provides a mechanism for multiplexing spatial with nonspatial information, such as task specific memory items (Wood et al., 2000) and sensory inputs from the lateral entorhinal cortex (Rennó-Costa et al., 2010). The independence of each neuron naturally explains the robustness of phase precession against intrahippocampal perturbations (Zugaro et al., 2005), an observation which is difficult to reconcile with models based on assembly interactions. Depending on the exact nature of the single cell phase code, we have shown that independent phase coding can enable theta sequences to be maintained with arbitrary global remapping. This flexibility may maximize the number and diversity of spatial representations that CA1 can provide to downstream structures.

Independent phase coding leads to several new and experimentally testable predictions that distinguish mechanisms of CA1 function during theta states. Firstly, an absence of coordination within or between assemblies has the advantage that neural interactions do not interfere with sequence generation after global remapping. Rather, for independent coding models the constraints on sequence generation following remapping arise from the nature of the phase code. With linear phase coding the set of sequences available to the network is fixed, resulting in a limited set of place field configurations with a particular mathematical structure (Figure 7). Interestingly, the remappings observed in grid modules (Fyhn et al., 2007), but not CA1, are consistent with those predicted for networks with a single fixed pacemaker input which forms a phase chart. More complex pacemaker systems with multiple charts could explain CA1 population activity during theta oscillations and “preplay”, which suggests a limited remapping capacity for CA1 (Dragoi and Tonegawa, 2011, 2013). Alternatively, sigmoidal phase coding massively increases the flexibility for global remapping as cells can remap arbitrarily while maintaining coherent theta sequences within each spatial representation (Figure 8). Secondly, linear and sigmoidal phase coding predict distinct MPO dynamics. With linear phase coding the temporal frequency of each MPO is independent of the animal’s location. With sigmoidal phase coding, the MPO frequency increases inside the place field, so that phase precession precession occurs inside but not outside the place field. In this case, only the spiking assembly behaves as a traveling wave, whereas the MPOs of cells with place fields distant from the animal are phaselocked to the LFP. Sigmoidal phase precession could emerge due to inputs from upstream structures (Chance, 2012) or be generated intrinsically in CA1 place cells (Leung, 2011). Finally, in contrast to linear phase coding populations, sigmoidal phase coding populations do not require additional directional information to generate directed theta sequences in open environments. Instead, sigmoidal theta sequences are determined solely by the recent trajectory of the rat through the set of place fields. This is consistent with recent observations of reversed theta sequences during backwards travel (Cei et al., 2014).

Independent coding implies that the spiking activity of CA1 place cell populations shows no correlations beyond that generated by a fixed rate and theta phase code in each cell. In other words, while the mutual dependence of each cell on LFP theta phase and location induces strong signal correlations, there are no additional correlations in the activity of the network. Because CA3 neurons immediately upstream from CA1 are connected by dense recurrent collaterals (Miles and Wong, 1986; Le Duigou et al., 2014), there are likely to be substantial additional correlations in the input to CA1, which might be expected to induce deviations from the independent population code outlined here. However, feedback inhibition motifs such as those found in CA1 are known to be able to counteract such correlations (Renart et al., 2010; Bernacchia and Wang, 2013; Tetzlaff et al., 2012; King et al., 2013). Hence, we suggest that the local inhibitory circuitry in CA1 removes any additional correlations present in its input in order to generate an independent population code. A major advantage of such a decorrelated representation is that it provides a highly readable and information rich code for working and episodic memory in downstream neocortex. In particular, a downstream decoder with access to an independent population code need only extract the stereotyped correlational patterns associated with traveling waves under a given place field mapping, allowing it to flexibly decode activity across a large number of spatial representations. Decoding in the presence of additional correlations would likely lead to a loss of information. While this loss can to some extent be limited by including knowledge of these additional correlations (Nirenberg and Latham, 2003; Eyherabide and Samengo, 2013), this likely requires a high level of specificity and therefore a lack of flexibility in the decoder. The flexibility afforded by an independent population code may therefore provide an optimal format for the representation and storage of the vast number of spatial experiences and associations required to inform decision making and guide behavior.

## EXPERIMENTAL PROCEDURES

We simulated data from a population of place cells with place field centers *x_c_* and width *σ* which precess linearly through a phase range of Δ*ϕ* over a distance 2*R* on a linear track using Equation (A3.6). To model discrete place cells with discrete action potentials, the firing rate *r* was binned over *x_c_* and time *t* respectively. The initial phase *ψ_s_* was either taken as 0, or a uniform random variable *ψ_s_ ∈* [0, 2*π*) set at the beginning of each run. In all simulations, parameters were set as: 2*R* = 37.5 cm (Maurer et al., 2006), Δ*ϕ* = 2*π*, *σ* = 9 cm, *f_θ_* = 8 Hz, *N*_spikes_ = 15. All simulations were performed using Matlab 2010b. To replicate the results of Harris et al. (2003), we simulated constant speed movement along a linear track, using a mean speed of 35 cm/s and standard deviation of 15 cm/s. We simulated motion in each direction, using the same set of place fields in each case. We estimated the preferred firing phase at each location from the simulated data using the methods stated in Harris et al. (2003), using either single-direction data or data consisting of runs in both directions to generate different phase fields. The prediction analyses were performed according to the methods given in Harris et al. (2003). In Figure 4C, the optimal prediction timescale for each phase locking value was chosen. This was done separately for the peer only case and the peer plus phase field case (other cases do not involve a peer prediction timescale).

To compare the sequence path length in spiking data generated from the traveling wave model to experimental data, we followed the decoding methods outlined in Maurer et al. (2011). Briefly, this involves constructing trial averaged time by space population activity matrices and decoding the location represented by the population in each time bin over the theta cycle. The decoded path length is measured as the largest distance between decoded locations within the theta cycle. To test the influence of phase locking in this analysis, *k* was varied incrementally from 0 to 6 and the sequence path length for the resulting data was calculated in each case. We used the same spatial and temporal bins (0.7 cm and 20*◦* of LFP *θ*) as the original study. To calculate the fast and slow slopes, we generated the contour density plots described by Maurer et al. (2011) using the same parameters as the sequence path length analysis. We simulated 100 trials for each running speed. We then divided these 100 trials into 10 subsets of 10 and applied the contour analysis to each subset. We fitted the fast slope to the 95% contour of the central theta peak, and measured the slow slope as the line joining the maximum of the top and bottom peaks of the central 3. We averaged over the results from each subset to obtain the final value. The analytical value for the fast slope in the limit of high phase locking is *FS* = *v_p_/*(360*f_θ_*), where the denominator arises due to the normalization to cm/deg in the analysis of (Maurer et al., 2011). Similarly for zero phase locking, *FS* = *v/*(360*f_θ_*). The analytical value for the slow slope is independent of phase locking *SS* = *v/*(360*f_θ_*). Upper and lower bounds for the slow slope were therefore fitted assuming the reported running speed is accurate, and that the LFP theta frequency is in the range 4 Hz < *f*_*θ*_ < 12 Hz.

To reproduce the results of Gupta et al. (2012), we used the significant sequence testing protocol and Bayesian decoding algorithm described therein, with spatial binning set as 3.5 cm, as in the original study. Briefly, the significant sequence testing analysis tests if population activity within a theta cycle has significant sequential structure, whereas the Bayesian decoding algorithm decodes the ahead and behind lengths encoded in the theta cycle. For Figure 6A, we varied phase locking and running speed independently and generated spiking data for each pair of values. We then applied the significant sequence detection methods for each resulting data set to obtain the fraction of significant sequences in each case. For Figure 6B&C, we set *k* = 0. We generated 160,000 theta cycles, each with a running speed drawn from a normal distribution with mean 30 cm/s and standard deviation 15 cm/s and then discarded speeds less than 10 cm/s. We applied the significant sequence detection algorithm to this data before applying the Bayesian decoding algorithm to the significant sequences to find the sequence path lengths, ahead lengths and behind lengths.

To reproduce the results of Foster and Wilson (2007), we generated data from 1000 theta cycles, each with a running speed drawn from the same distribution as for the Bayesian decoding analysis. Following the protocol outlined by Foster and Wilson (2007), we found the set of all spike phases for each cell when the rat was at each position and analyzed events defined as 40 ms windows around the peak firing rate (i.e., LFP theta trough). For the shuffling analysis, spikes in each event were replaced by another spike taken from the same position and cell. The new spike time was then calculated by interpolation between the closest two LFP theta troughs of the original spike, as reported in the original study. 100 such shuffles were performed for each event, and the correlations between cell rank order and time were calculated in each case.

## ACKNOWLEDGEMENTS

This work was supported by the EPSRC, BBSRC and MRC. We thank Kamran Diba and Iris Oren for helpful comments on the manuscript.

